# Spatial metabolomics reveals localized impact of influenza virus infection on the lung tissue metabolome

**DOI:** 10.1101/2021.11.22.469643

**Authors:** Danya A. Dean, London Klechka, Ekram Hossain, Adwaita R. Parab, Krystin Eaton, Myron Hinsdale, Laura-Isobel McCall

## Abstract

The influenza virus (IAV) is a major cause of respiratory disease, with significant infection increases in pandemic years. Vaccines are a mainstay of IAV prevention, but are complicated by consideration of IAV’s vast strain diversity, manufacturing and vaccine uptake limitations. While antivirals may be used for treatment of IAV, they are most effective in early stages of the infection and several virus strains have become drug resistant. Therefore, there is a need for advances in IAV treatment, especially host-directed, personalized therapeutics.Given the spatial dynamics of IAV infection and the relationship between viral spatial distribution and disease severity, a spatial approach is necessary to expand our understanding of IAV pathogenesis. We used spatial metabolomics to address this issue. Spatial metabolomics combines liquid chromatography-tandem mass spectrometry of metabolites extracted from systematic organ sections, 3D models and computational techniques, to develop spatial models of metabolite location and their role in organ function and disease pathogenesis. In this project, we analyzed plasma and systematically sectioned lung tissue samples from uninfected or infected mice. Spatial mapping of sites of metabolic perturbations revealed significantly lower metabolic perturbation in the trachea compared to other lung tissue sites. Using random forest machine learning, we identified metabolites that responded differently in each lung position based on infection, including specific amino acids, lipids and lipid-like molecules, and nucleosides. These results support the implementation of spatial metabolomics to understand metabolic changes upon IAV infection and to identify candidate pathways to be targeted for IAV therapeutics.

**Importance:** The influenza virus is a major health concern. Over 1 billion people become infected annually despite the wide distribution of vaccines, and antiviral agents are insufficient to address current clinical needs. In this study, we used spatial metabolomics to understand changes in the lung and plasma metabolome of mice infected with influenza A virus, compared to uninfected controls. We determined metabolites altered by infection in specific lung tissue sites and distinguished metabolites perturbed by infection between lung tissue and plasma samples. Our findings highlight the importance of a spatial approach to understanding the intersection between lung metabolome, viral infection and disease severity. Ultimately, this approach will expand our understanding of respiratory disease pathogenesis and guide the development of novel host-directed therapeutics.

## Introduction

Influenza virus outbreaks are a continuous public health issue. Seasonal global epidemics caused by both influenza A viruses (IAV) and influenza B viruses cause 300,000-500,000 deaths each year (1). Vaccinations are the current method of prevention, but they fail to account for every possible viral strain. Antiviral drugs are used for treatment of IAV but are most effective within a short window during early infection. Additionally, it is believed that some strains have developed resistance to these drugs (2). One study indicated that while influenza A(H1N1)pdm09-infected intensive care unit patients treated with neuraminidase inhibitors have greater survival rates than untreated patients, one in four treated patients still die (3). These findings indicate a strong need for new treatments for IAV infection, and the potential for host-targeted therapeutics to supplement antiviral agents. Their development, however, necessitates an understanding of disease pathogenesis, which remains incompletely elucidated for IAV.

We used metabolomics to identify and analyze metabolites affected by IAV infection. Metabolomics is a method of analysis that focuses on small molecules involved in biological processes. This technique allows us to gain insight on the host chemical response to viral infection. A comprehensive understanding of the relationship between host and virus could aid in the development of more effective prevention and treatment options. Previous studies applied metabolomic methods to lung tissue and serum during IAV infection. These studies found that nucleosides such as uridine, lipids such as sphingosine, sphinganine, and amino acid metabolites such as kynurenine are increased during infection in lung tissue (4). Additionally, carbohydrates such as mannitol, myo-inositol and glyceric acid are decreased during active infection. However, location of IAV within the respiratory tract is dynamic. Viral localization and location of tissue damage within the respiratory tract also influences disease symptoms, disease severity and transmissibility of the infection (5, 6, 7, 8). Thus, a spatial perspective is necessary with regards to IAV and tissue metabolism. Chemical cartography is an approach that combines liquid chromatography-mass spectrometry (LC-MS) with 3D visualizations, leading to detailed spatial maps of metabolite distribution compared to pathogen load, tissue damage, or immune responses (9, 10, 11). This approach, when applied to other infectious diseases enabled the discovery of new treatments for these conditions (10). This method has been used to study the impact of cystic fibrosis on the local lung metabolome (12, 13), but had not previously been applied to IAV infection.

We therefore hypothesized that spatial metabolomics could provide new insights regarding IAV infection. Using this method, we analyzed the distribution of small (*m/z* 100-1,500) metabolites within the infected trachea and lung, in comparison to plasma samples and to uninfected animals. We identified changes in the lung metabolome and determined limited overlap in metabolites perturbed by IAV infection between lung tissue and plasma. Additionally, we identified several metabolites altered by infection such as amino acids, lipids and lipid-like molecules, and nucleosides. Interestingly, these metabolites were differentially affected at each lung position. Ultimately, our study highlights the application of spatial metabolomics to understand IAV infection and further our understanding of respiratory disease pathogenesis, to guide the development of novel host-directed therapeutics.

## Results

Viral distribution influences IAV transmissibility, viral reassortment and disease severity (5, 14, 15). While a few studies have investigated the changes in the metabolome during IAV infection, a spatial perspective of the metabolic disturbances is lacking (4, 16–18). We therefore used spatial metabolomics to identify candidate pathways and metabolites altered by infection in specific lung locations. Plasma, trachea and lungs were collected from IAV-infected mice at 3 days post-infection. Lungs were systematically sectioned into 11 segments (Fig. 1A), and all samples analyzed by liquid chromatography-tandem mass spectrometry (LC-MS/MS), followed by 3D reconstruction of metabolomics data. In addition, tissue homogenate bioluminescence was measured as an indicator of local viral burden in each tissue segment.

**Figure 1.**
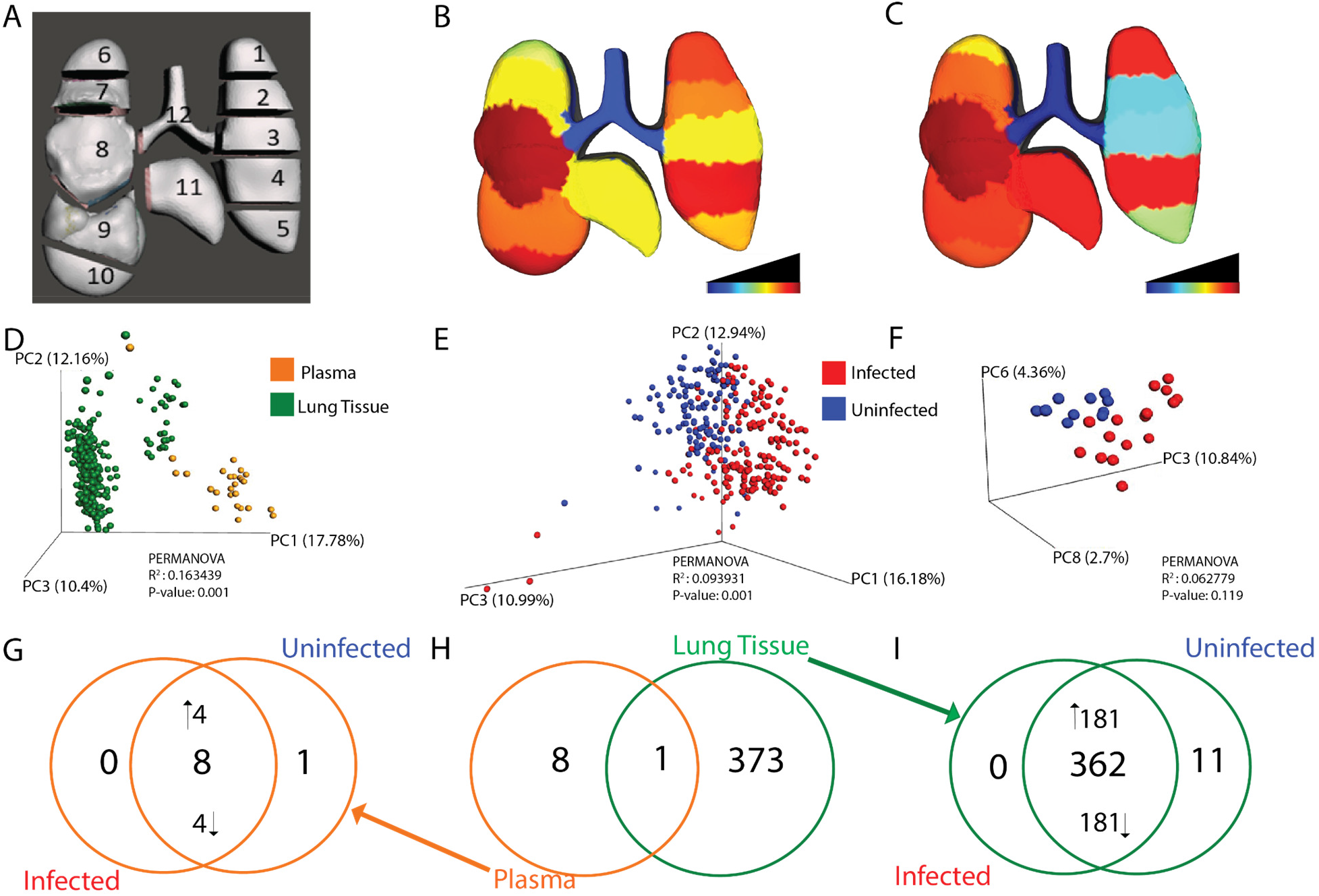
Localized impact of IAV infection in lung tissue and plasma. **A:** 3D model of lung tissue showing sampling positions. **B:** Median viral burden distribution of IAV. Lower infection levels were observed in the trachea (Dunn’s test, FDR-corrected, p<0.05 for comparisons between trachea and all positions except left lung middle (position 3)). **C:** Magnitude of metabolic perturbation compared to uninfected (PERMANOVA R^2^). **D:** Principal coordinate analysis (PCoA) plot showing differences in overall metabolome by sample type (PERMANOVA p-value <0.001). **E:** Impact of infection on overall lung tissue metabolome (PERMANOVA p-value <0.001). **F:** No significant impact of infection on the overall plasma metabolome by PCoA. **G-I:** Unique and common metabolites perturbed by infection **G:** Most metabolites perturbed by infection in plasma samples are found in both infected and uninfected samples, albeit at different levels. 4/8 metabolites were increased by infection and 4/8 were decreased by infection, while only one of the statistically significant metabolites was uniquely observed only in uninfected plasma samples. **H:** Overlap of metabolites perturbed by infection in plasma (orange) or lung tissue all positions combined (green). **I:** Most metabolites perturbed by infection in lung samples are found in both infected and uninfected samples, albeit at different levels (181/382 metabolites increased by infection and 181/362 decreased by infection, with 11 of the statistically significant metabolites uniquely detected in uninfected lung tissue).

Using principal coordinate analysis (PCoA), we first compared lung and plasma metabolomes and found that the local lung tissue metabolome does not reflect the circulating metabolome (PERMANOVA <0.001, Fig 1D). Furthermore, lung tissue overall was found to be greatly impacted by infection (PERMANOVA <0.001, R^2^ 0.093931) (Fig 1E). A similar but non-significant, trend was seen between infected and uninfected plasma samples (Fig 1F).

Spatial analyses of lung tissue showed that viral load was largely localized to the lung tissue (positions 1-11) with minimal viral load in the trachea (position 12) (Dunn’s test p<0.05, FDR-corrected, for comparisons between trachea and all positions except left lung middle (position 3, Fig 1B)). Overall local impact of infection on the metabolome was quantified using PERMANOVA R^2^ at each sampling site. Magnitude of metabolic perturbation was variable between tissue segments, with the highest degree of metabolic perturbation in two segments of the left lung (position 1 and position 4), and the right lung middle lobe (position 8), whereas the trachea metabolome was least affected (R^2^ range from 0.05 in trachea to 0.18 at position 4; Fig 1C and Table S15). Thus, sites of highest and lowest metabolic perturbation match sites with high vs low viral load, respectively, enabling insight into the specific metabolites most affected by infection.

We next sought to determine whether the specific metabolites perturbed by infection differed between lung and plasma (Fig 1G-I). We used a random forest classifier, applied to plasma on the one hand, and to lung tissue on the other hand (all positions combined). After applying significance cutoffs (see Materials and Methods), this approach yielded a total list of 9 plasma metabolites and 374 lung tissue metabolites significantly perturbed by infection. There was strikingly limited overlap of infection-perturbed metabolites between lung tissue samples and plasma samples, indicating that both sites respond differentially to infection (Fig 1H) and concurring with overall PCoA analysis findings (Fig 1D). We then sought to assess whether the metabolites perturbed by infection were uniquely elicited by infection, or found under both conditions but at differential levels. The majority of infection-perturbed metabolites were common to infected and uninfected samples but found at different levels, with a minority uniquely detected in uninfected samples only (Fig 1I). A similar trend was seen in infected and uninfected plasma samples, where most infection-perturbed metabolites were present in both infected and uninfected samples, albeit at different levels (Fig 1G). This signifies that the effect of IAV infection on the metabolome is primarily on metabolite levels rather than induction of novel metabolites.

Random forest classifier was also used to analyze the common and unique responses of each lung and tracheal tissue site to infection. A random forest model was built for each lung position and for plasma, classifying infected versus uninfected samples. Strikingly, most infection-impacted metabolites were only affected at one or a few tissue sites (Fig 2A). This is not due to divergence in overall metabolome between sites, as analysis of all metabolites, irrespective of abundance, revealed large commonality across lung tissue sites and plasma (Fig 2B). Likewise, infection-perturbed metabolites do not overlap appreciably with metabolites differing in abundance between lung and plasma (Fig S3). Very few perturbed metabolites were identified for position 12 (trachea), perhaps as a consequence of the low viral load at that site (Fig 1B).

**Figure 2:**
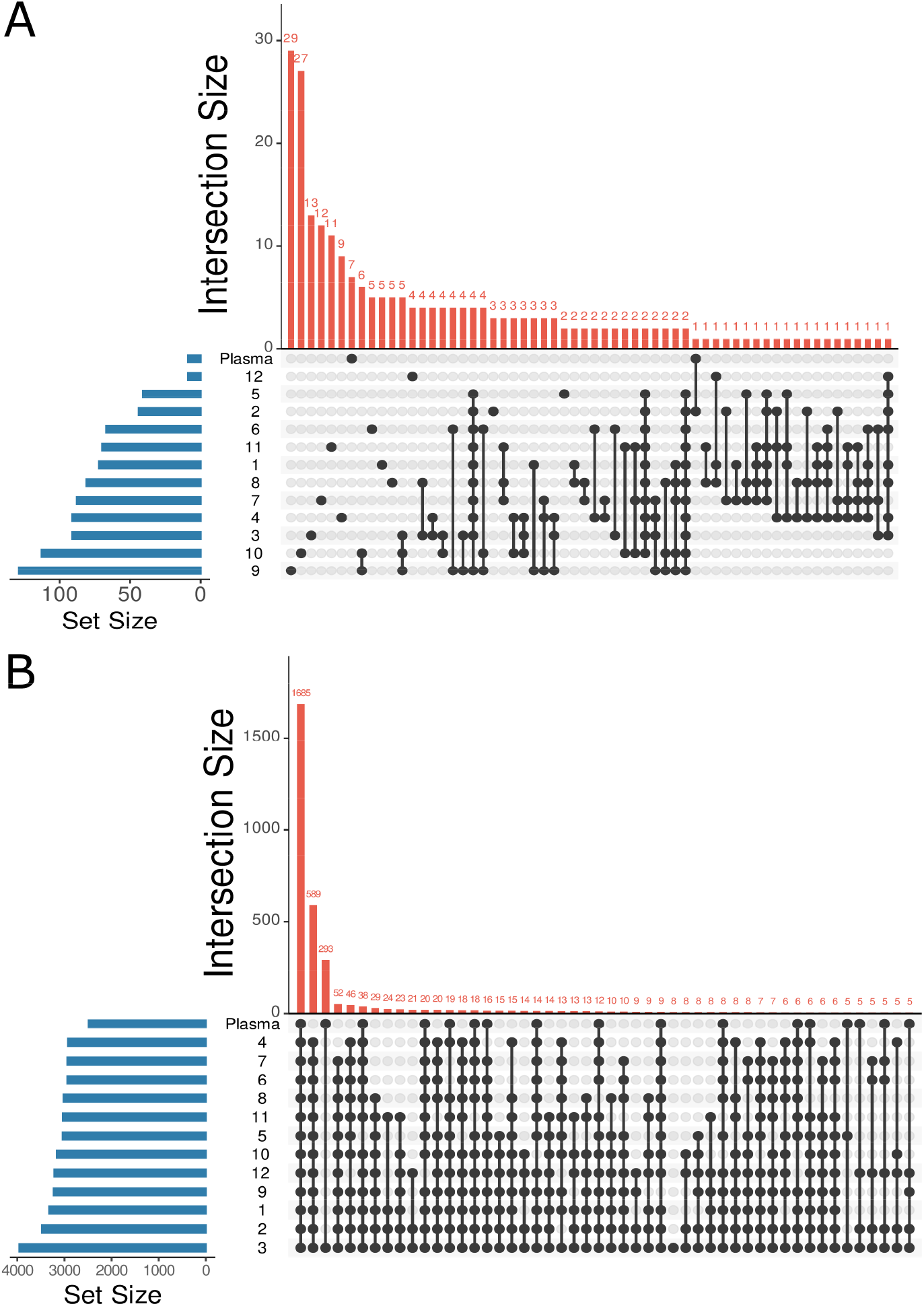
Location-specific impact of IAV infection on metabolism. A: UpSet plot (Number of intersections to show: 60) showing uniqueness of statistically significant infection-impacted metabolites in lung tissue and plasma perturbed, based on 13 random forest models analyzing each lung segment and plasma separately. B: UpSet plot (Number of intersections to show: 50) of total metabolites found in lung tissue and plasma showing large commonality across tissue sites and plasma. Position numbers as in Fig. 1A.

Metabolites perturbed by infection were annotated using molecular networking (19). Among these, carnitine, glutamine, kynurenine, and cytosine were found to be significantly and markedly perturbed by IAV infection and in a spatial manner (Fig 3, Fig S1). Glutamine, cytosine, and kynurenine were all significantly increased by IAV infection at all tissue sites except the trachea (Wilcoxon FDR-corrected p-value <0.05 at all sites except trachea) (Fig 3, Fig S1). The opposite trend was seen for carnitine which was decreased by infection at all tissue sites except the trachea (Wilcoxon FDR-corrected p-value <0.05 at all sites except trachea) (Fig 3, Fig S1). In contrast, these metabolites were not significantly affected by infection in the plasma.

**Figure 3:**
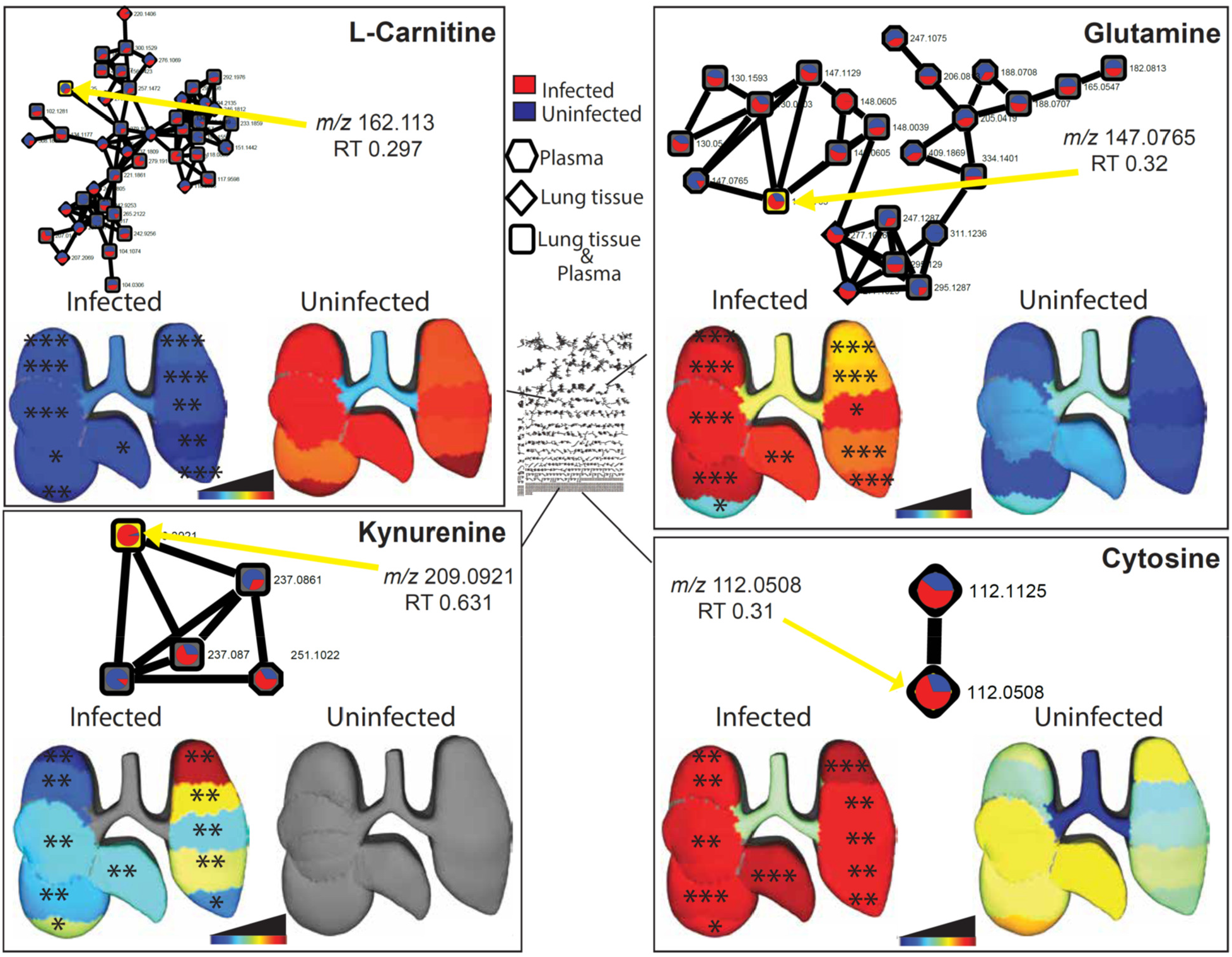
Representative metabolites perturbed by IAV infection. Molecular networks display summed peak area of metabolites in infected (red) and uninfected (blue) samples. Each connected node represents structurally-related metabolites, as determined by molecular networking. 3D lung ‘ili plots show median peak area of each displayed metabolite across tissue sites. Wilcoxon FDR-corrected p-values comparing matched infected and uninfected lung tissue sites *<0.05; **<0.01; *** <0.001.

Additional metabolites perturbed by infection in plasma and lung tissue include amino acids, acylcarnitines and nucleobases (Fig 4, Table S1, Table S2), with contrasting effects between lung tissue positions and sample types. For example, amino acids had dissimilar effects following IAV infection in plasma and lung tissue samples (Fig 4): methionine was decreased in infected plasma samples (Wilcoxon FDR-corrected p-value <0.001) while phenylalanine and L-citrulline were increased in infected lung tissue samples (Wilcoxon FDR-corrected p-value <0.05) (Fig 4 A, E-F). However, the dipeptide methionine-phenylalanine was decreased in infected lung tissue (Wilcoxon FDR-corrected p-value <0.05) (Fig 4K).Acylcarnitines (oleylcarnitine and palmitoylcarnitine) were significantly increased in infected plasma samples (Wilcoxon FDR-corrected p-value <0.001) compared to uninfected samples (Fig 4B-C). Guanine, S-adenosy-homocyteine, spermidine, 4-hydroxynonenal, and hemin cation were all significantly increased upon infection (Wilcoxon FDR-corrected p-value <0.05) at select lung tissue sites (Fig 4F-H, G and L). Infection had the opposite effect on palmitamide, which was significantly decreased in infected tissue samples (Fig 4I). In contrast, none of these metabolites apart from phenylalanine were significantly affected by infection in the trachea.

**Fig 4:**
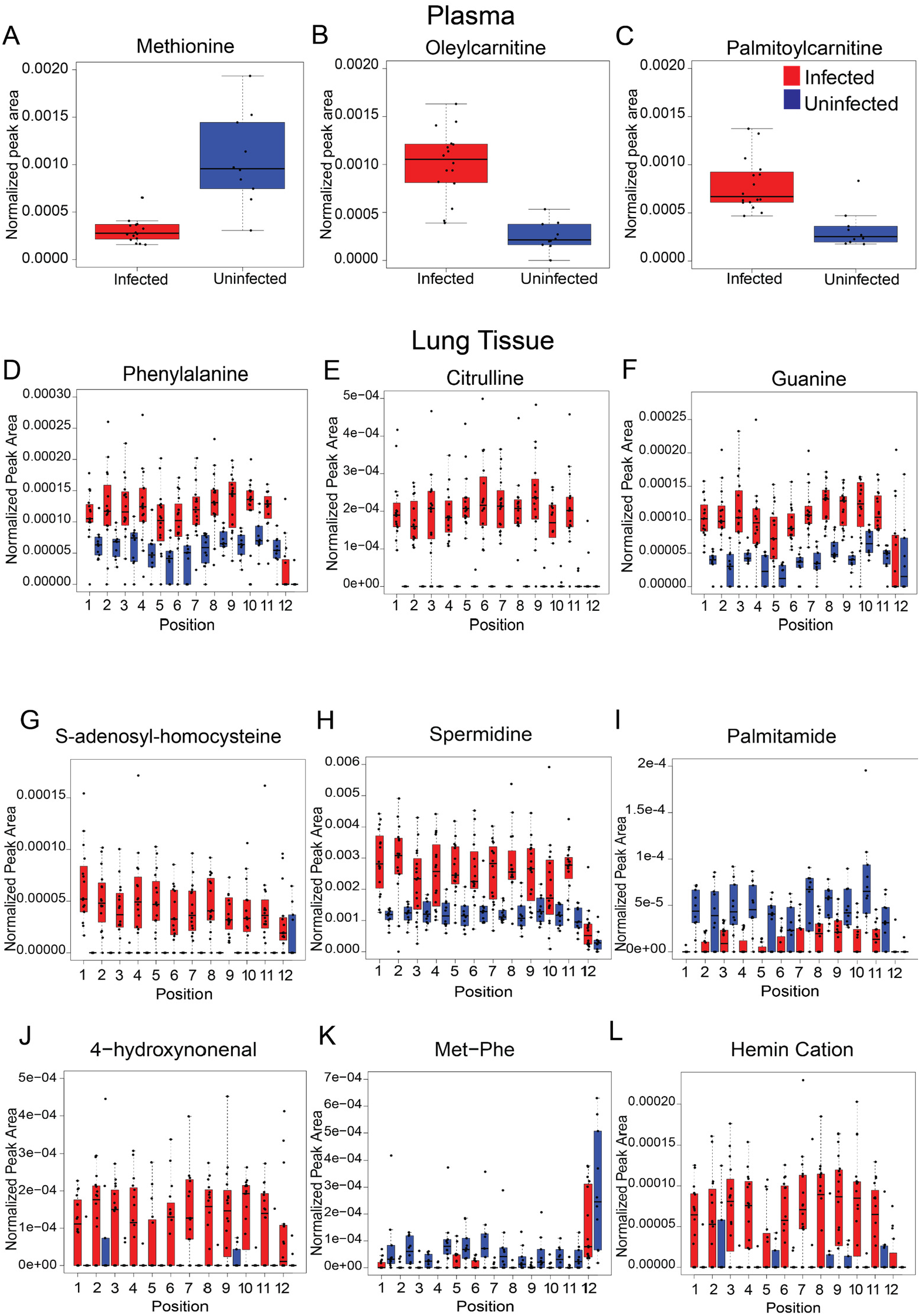
Representative metabolites perturbed by IAV infection in plasma and lung tissue. **A-C:** Statistically significant metabolites altered by infection in plasma (Wilcoxon FDR-corrected p-value <0.001). **A:** Methionine (*m/z* 150.058, RT 0.323 min). **B:** Oleyl L-carnitine (*m/z* 426.357, RT 2.982 min). **C:** Palmitoylcarnitine (*m/z* 400.341, RT 2.952 min). **D-L:** Statistically significant metabolites altered by infection in lung tissue. **D:** Phenylalanine (*m/z* 166.042, RT 2.246 min) statistically different at all positions (Wilcoxon FDR-Corrected p-value <0.05) except positions 2, 5, 10 and 12. **E:** Citrulline (*m/z* 198.085, RT 0.309 min) statistically significant at all positions (Wilcoxon FDR-Corrected p-value <0.05) except position 12. **F:** Guanine (*m/z* 150.977, RT 0.291 min) statistically different at all positions (Wilcoxon FDR-Corrected p-value <0.05) except position 12. **G:** S-Adenosyl-homocysteine (*m/z* 385.212, RT 2.166 min) statistically significant at all positions (Wilcoxon FDR-Corrected p-value <0.05) except position 12. **H:** Spermidine (*m/z* 146.165, RT 0.287 min) statistically significant at all positions (Wilcoxon FDR-Corrected p-value <0.05) except positions 3, 10, and 12. **I:** Palmitamide (*m/z* 256.073 RT 2.563 min) statistically significant at all positions (Wilcoxon FDR-Corrected p-value <0.05) except positions 2, 5, 6, 7, 11, and 12. **J:** 4-Hydroxynonenal (*m/z* 139.112, RT 2.644 min) statistically significant at all positions (Wilcoxon FDR-Corrected p-value <0.05) except positions 1, 2, 3, 5, 11, and 12. **K:** Met-Phe (*m/z* 297.123, RT 2.444 min) statistically significant at all positions (Wilcoxon FDR-Corrected p-value <0.05) except positions 3, 4, 6, and 7. L: Hemin cation (*m/z* 616.358, RT 2.994 min) statistically significant at all positions (Wilcoxon FDR-Corrected p-value <0.05) except positions 3, 4, 9, and 10.

## Discussion

Here, we generated spatial maps of the metabolic impact of IAV infection on the mouse lung and trachea. This approach revealed differential effects of infection across tissue sites and between lung and plasma, as well as differential viral burden between trachea and lung tissue. Based on random forest analysis and molecular networking, infection-impacted metabolites of biological significance were annotated as acylcarnitines, amino acids, phospholipids, and nucleotides, amongst others.

L-carnitine was significantly decreased by IAV infection at all lung positions except the trachea (Fig 3 top left). Likewise, short chain acylcarnitines such as CAR 2:0 and CAR 6:0 were also decreased in lung tissue (Table S3-14). In contrast, in both plasma and lung tissue, different long chain acylcarnitines were increased by infection, including oleyl carnitine (CAR 18:1) and palmitoylcarnitine (CAR 16:0) in plasma and CAR 14:1 and CAR 20:1 in lung tissue (Fig 4B-C, Table S1-14). Carnitines and acylcarnitines are key intermediates in energy production via fatty acid beta oxidation, and influenza virus replication is sensitive to fatty acid beta oxidation activity (20),(21). Likewise, these metabolic alterations may also be contributing to the differential responses to vaccination in obese vs lean animals (22).

Our study indicated that glutamine is increased upon IAV infection (Fig 3 top right). T cell proliferation and cytokine secretion relies heavily on glutamine presence (23). Thus, elevated glutamine levels may enable anti-viral immunity IAV-infected cells are also more dependent on glutamine availability than uninfected cells for survival, suggesting a pro-survival effect of our observed elevated glutamine levels (24). Phenylalanine and citrulline were increased in infected lung tissue, in contrast with methionine, which was decreased in plasma (Fig 4D-E). Both phenylalanine and citrulline are involved in immune response by aiding in T-cell function (25). Phenylalanine aids in activation of T-cells while citrulline has downstream effects through the production of arginine which is used for T-cell growth and response (25–27). Upregulation of both amino acids suggest their role in overall immune response to IAV.

Nucleotides, cytosine and guanine were elevated by infection in all lung sites except trachea (Fig 3 bottom left and Fig 4F). Pyrimidine nucleotide biosynthesis is elevated upon IAV infection, and indeed its replication was dependent on pyrimidine biosynthesis (28–30). Interestingly, cytosine was found discriminatory in the plasma between RT-PCR positive COVID patients and RT-PCR COVID negative patients (29).

Several of the metabolites annotated and identified as infection-impacted in the lung tissue and plasma in this study are congruent with prior studies of respiratory infection (4). Amino acids and nucleotides in particular were upregulated in lung tissue in other IAV studies as well as in *Mycobacterium tuberculosis* infection (TB) and respiratory syncytial virus (RSV) studies (4, 31–33). Phenylalanine was increased in mouse lung tissue in another IAV study and in the serum and lung of mice infected with TB, while L-citrulline is elevated in mouse lung tissue infected with RSV and in mice infected with TB (4, 17, 31, 32, 34). This coincides with our findings showing elevations of these amino acids in IAV-infected lung tissue (Fig 4D and E).

Glutamine was identified in mouse lung tissue infected with TB and was elevated alongside other immune response amino acids, corresponding with our study (Fig 3 top right) (32, 34). Kynurenine, an infection-induced anti-inflammatory molecule, is consistently upregulated in mouse lung tissue of several respiratory infection studies including TB, RSV, and our IAV study (Fig 3 bottom left) (4, 31, 33, 34). Upregulation in cytosine in the lung tissue, oleylcarnitine in plasma, and several phospholipids in the lung tissue are also consistent with current literature (Fig 3 bottom right, Fig 4B and Table S1) (4, 17, 33).

Analysis of circulating metabolites has been performed in multiple studies on IAV vaccinology (35), (36). Our findings of differential impact of infection on lung and plasma metabolites (Fig. 1H, Table S1, Table S2) indicate that the metabolic changes observed in those studies may not be directly linked to lung metabolic patterns, and this discrepancy was not due purely to differences between plasma and lung overall (Fig. 2B). Likewise, the majority of COVID-19 metabolomic studies have relied on serum or plasma samples (37). By extension, based on this study’s results, they may not be relevant to the pathogenesis of COVID-19 in the lung, hampering translatability for drug development purposes. Thus, studies seeking to build on metabolomics to design new treatments should rely on analysis of the affected organ, rather than biofluids, even if the latter are more readily available.

We also observed differential impact of infection across lung sections, with the majority of infection-perturbed metabolites only significantly perturbed at one lung segment (Fig. 2A). The trachea in particular was especially divergent from the lung lobes in terms of viral burden (Fig. 1B), overall magnitude of infection-induced metabolic perturbations (Fig. 1C) and specific metabolic changes (Fig. 3, Fig. 4). Jointly, these results highlight the strength of our spatial perspective.

As with any untargeted metabolomics studies, a significant fraction of infection-impacted metabolites could not be annotated (∼65%). The most commonly observed metabolite subclasses in our dataset overall were amino acids, peptides, and analogues, glycerophosphocholines, amines and fatty amides. Although this represents a broad diversity of metabolite classes, nevertheless complementary metabolite extraction or data acquisition methods could further expand this list. We further acknowledge that the mouse model of influenza virus infection and mouse-adapted influenza viral strain may not be the most representative of the functional lung alterations that would occur during human infection (38). Indeed, the predominance of lower respiratory tract metabolic alterations over changes in the trachea observed in this study. may be a consequence of this model. A further limitation is that we focused on a single time point, in a lethal infection model. However, our findings serve as proof-of-concept of the applicability of our approach to study respiratory viruses, with future work applying this method in other biological systems.

We anticipate our findings to serve as a resource upon which the research community can build to study the impact of different disease modifiers on the relationship between spatial changes in the lung metabolome and disease severity, for example the impact of age, comorbidities or treatment. Our findings and our approach also serve as a framework to study how the metabolome is restored in a spatially-dependent fashion during recovery from respiratory infection, or fails to recover during chronic disease in additional disease models, and to identify markers of treatment response and infection outcome. Overall, we anticipate our approach to be broadly applicable to many other respiratory infections, helping expand our understanding of respiratory disease pathogenesis and to drive the development of host-targeted therapeutic regimens.

## Materials and Methods

### *In vivo* infection

All vertebrate animal studies were performed under a protocol approved by Oklahoma State University Institutional Animal Care and Use Committee (protocol number VM20-36), in accordance with the USDA Animal Welfare Act and the Guide for the Care and Use of Laboratory Animals of the National Institutes of Health.

Female C57BL/6J mice were anesthetized with ketamine and xylazine intraperitoneally and then intranasally inoculated with PR8-Glu (39) (a generous gift from Dr. Peter Palese, Icahn School of Medicine at Mount Sinai, Department of Microbiology, New York, New York, USA) at 2 × 10^3^ pfu/mouse in 25-50 μl PBS. Controls were inoculated with PBS alone. Body weights were monitored daily. Three days later, mice were anesthetized with ketamine and xylazine and injected IV with a working solution of coelenterazine (GoldBio, St. Louis, Mo.) made according to standard protocol (40). Briefly, using a stock solution of 7.5 mg/ml in acidified alcohol, each mouse was administered 98 μg substrate in PBS (13 μl stock in 137 μl PBS) into the retroorbital sinus. After substrate administration, the thoracic cavity was opened followed by exsanguination by cardiac blood collection for serum and by lung tissue harvesting. Upper trachea, cranial to tracheal bifurcation, was also collected. Subsequently, samples were submerged in coelenterazine 0.3 mg/ml in a 96 well plate. Once collected the tissue pieces were blotted off and flash frozen in liquid nitrogen for later analyses.

### Metabolite extraction and UHPLC-MS/MS

A two-step metabolite extraction procedure was performed according to Want *et al*.(41) for both tissue and plasma samples. Samples were homogenized in LC-MS grade water with steel beads utilizing a Qiagen TissueLyzer at 25 Hz for 3 min, and 1 μl removed for luminescence analysis. Methanol was added for a final concentration of 50%, samples homogenized again for 3 mins and centrifuged at 16,000xg for 10 mins at 4ºC. The supernatant (aqueous extract) was collected, dried overnight in a Speedvac and frozen at −80ºC until LC-MS analysis. The pellet produced from centrifugation was collected for organic extraction via addition of 3:1 dichloromethane:methanol, homogenized for 5 mins and centrifuged for 10 mins at 4ºC. The organic extract was air dried overnight and then frozen at −80ºC until LC-MS analysis. Dried aqueous and organic extracts were resuspended in 1:1 methanol and water spiked with the internal standard sulfadimethoxine and combined. Samples were then sonicated, centrifuged, and the supernatant collected for analysis. A Thermo Scientific Vanquish UHPLC system was used for tissue and plasma analysis using a Kinetex 1.7μm C8 100 Å LC column (50 × 2.1mm). Chromatography was done with water + 0.1% formic acid (mobile phase A) and acetonitrile + 0.1% formic acid (mobile phase B), at a 0.5 mL/min flow rate (7.5 mins) with a 40°C column temperature. LC gradient can be found in Table 1. Data acquisition was performed in random sample order, with a blank and pooled quality control every 12 samples.To monitor instrumental drift, a 6-mix solution with 6 known molecules was run at the beginning and end of LC/MS/MS analysis. Calibration of the instrument was also done immediately prior to instrument analysis using Pierce LTQ Velos ESI positive ion calibration solution. MS/MS detection was conducted on a Q Exactive Plus (Thermo Scientific) high resolution mass spectrometer (Table 2). Ions were generated for MS/MS analysis in positive mode.

**Table 1:** LC Gradient

| Time(min) | Flow (ml/min) | %B | Curve |
| --- | --- | --- | --- |
| 0.000 | Run |  |  |
| 0.000 | 0.500 | 2.0 | 5 |
| 1.000 | 0.500 | 2.0 | 5 |
| 2.500 | 0.500 | 98.0 | 5 |
| 4.500 | 0.500 | 98.0 | 5 |
| 5.500 | 0.500 | 2.0 | 5 |
| 7.500 | 0.500 | 2.0 | 5 |
| 7.500 | Stop Run |  |  |

**Table 2:**
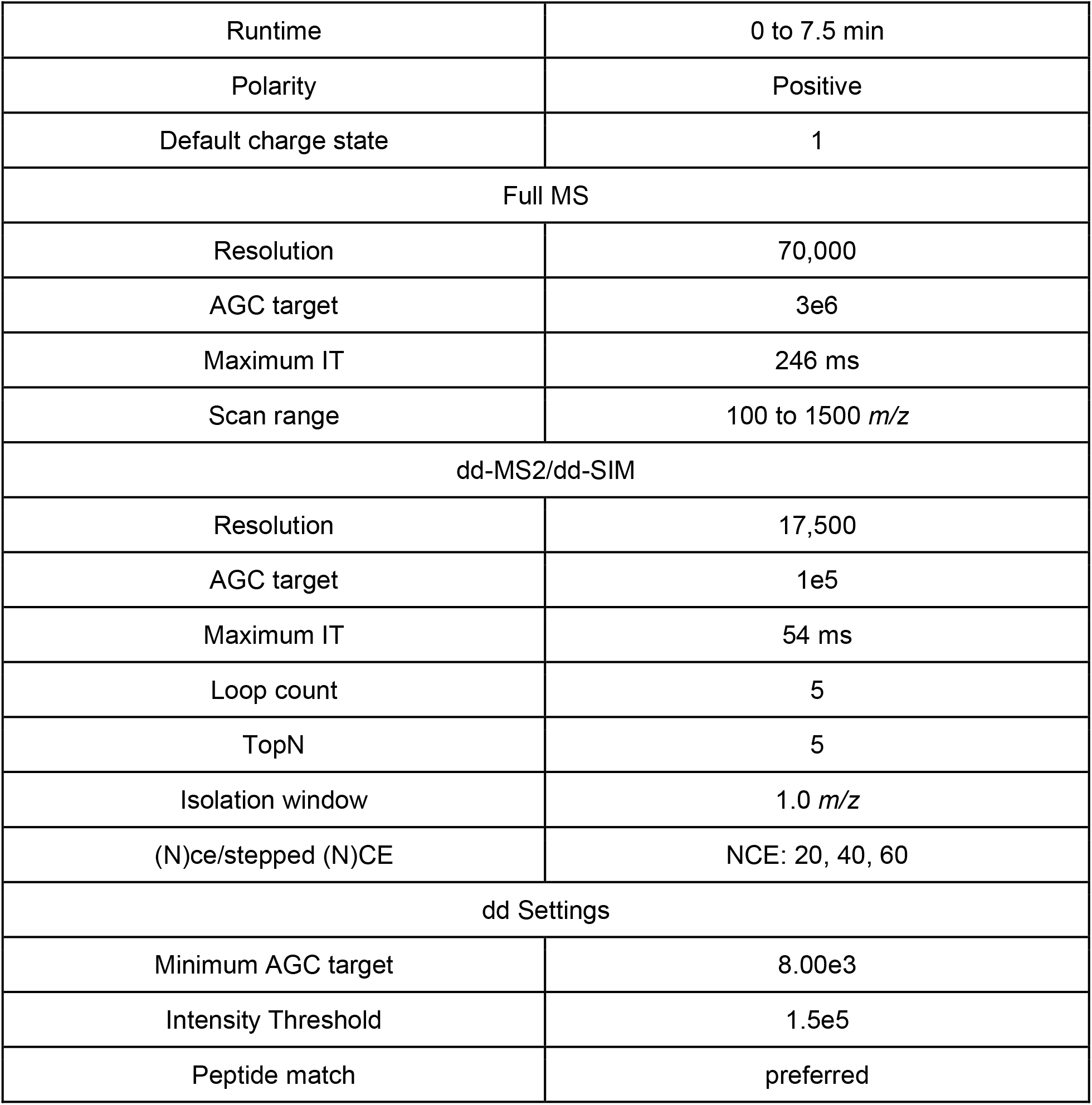
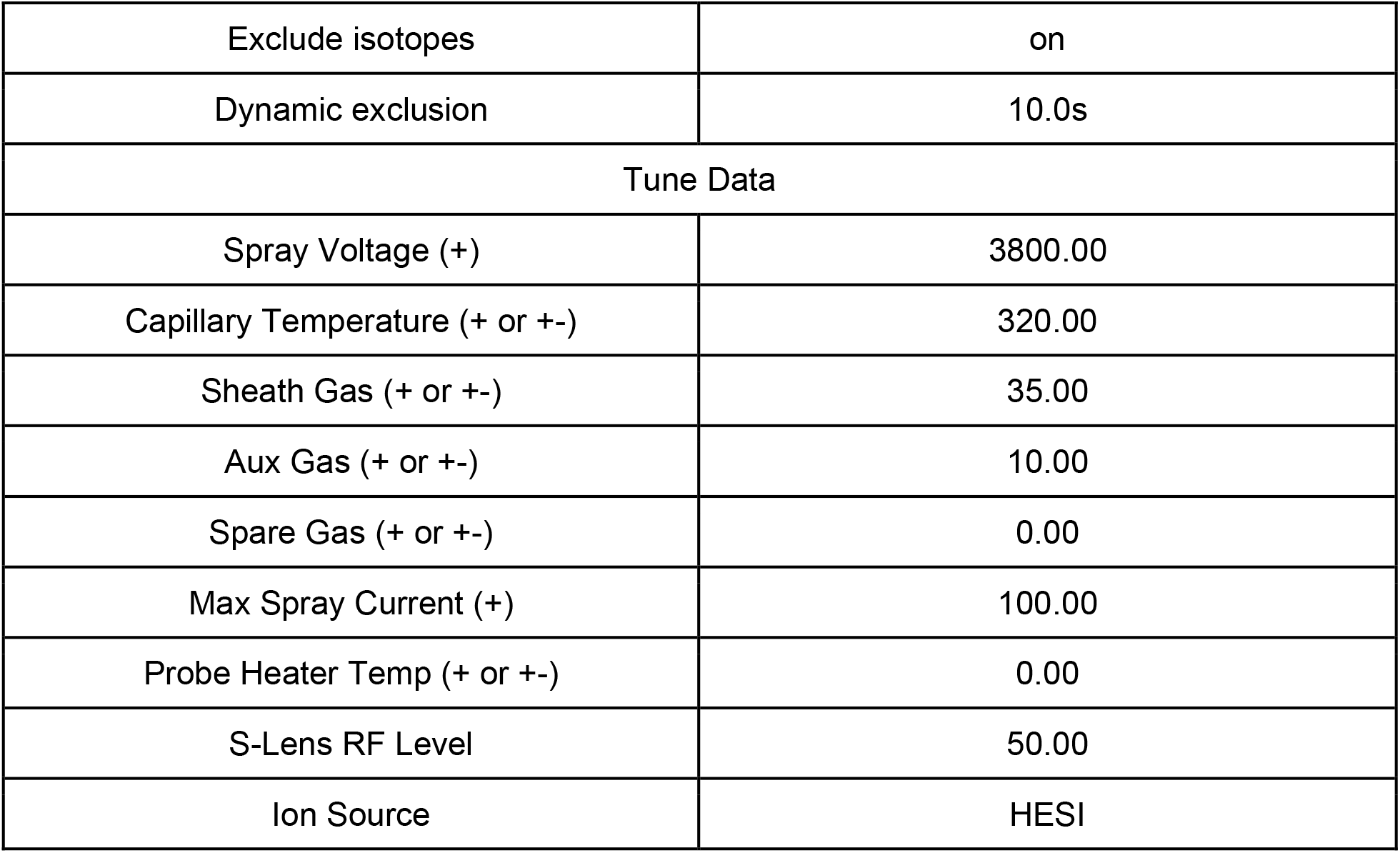
Q Exactive Plus (Thermo Scientific) instrument parameters

### Luminescence analysis

Coelenterazine was prepared according to GoldBio standard protocols. Briefly, 1 mg coelenterazine was added to 1 mL of acidified methanol to make a stock solution. The stock solution was made to a final concentration of 1.5 μM. A 1:10 solution of sample homogenate to colentrazine was analyzed on a GloMax Explorer (Promega).

### LC-MS Data analysis

Data analysis was performed using MZmine version 2.53, according to Table 3 parameters, to develop the feature table for further analysis. Blank removal with a 3-fold threshold was performed and Jupyter notebooks were used to perform total ion current (TIC) normalization. Principal coordinate analysis (PCoA) was performed on the TIC-normalized MS1 feature table using the Bray-Curtis dissimilarity metric in QIIME2 (42). Three-dimensional PCoA plots were developed and visualized using EMPeror (43). Lung three-dimensional model developed using Sketchup and Mesh lab, and modelling was completed using ‘ili (http://ili.embl.de/) (44).

**Table 3:** MzMine 2.53 parameters

| MS<sup>1</sup> |  |
| --- | --- |
| Retention Time (min) | 0-7.5 |
| Noise Level | 5E5 |
| MS<sup>2</sup> |  |
| Retention Time (min) | 0-7.5 |
| Noise Level | 1E3 |
| Chromatogram Builder |  |
| Mass list | Masses |
| Minimum Time span (min) | 0.01 |
| Minimum Height | 1.5E6 |
| m/z Tolerance (ppm) | 10.0 |
| Algorithm | Baseline cut-off |
| Chromatogram Deconvolution |  |
| *m/z* Range for MS<sup>2</sup> scan pairing (Da) | 0.001 |
| RT Range for MS<sup>2</sup> Scan pairing (min) | 0.2 |
| Minimum peak height | 1.5E6 |
| Peak duration range (min) | 0-3.5 |
| Baseline level | 5E5 |
| Deisotoping |  |
| *m/z* tolerance (ppm) | 10.0 |
| Retention time tolerance (min) | 0.5 |
| Monotonic shape | Yes |
| Maximum Charge | 3 |
| Representative Isotope | Most intense |
| Alignment |  |
| *m/z* Tolerance (ppm) | 10.0 |
| Weigh for *m/z* | 1 |
| Weight for Retention time | 1 |
| Retention time tolerance (min) | 0.5 |
| Row filtering |  |
| Minimum peaks in a row | 3 |
| Retention time (min) | 0.2-6.5 |
| Keep only peaks with MS<sup>2</sup> scans | YES |
| Reset Peak No. ID | YES |

Random forest analysis was conducted using R in Jupyter notebooks. Number of trees were restricted to 500 and random forest classifier cutoff was based on variable importance score of mean decrease accuracy >1. Lists were further restricted to FDR-corrected Mann-Whitney p-value less than 0.05 and fold change <0.05 or >2.0. Venn diagrams and UpSet plots were developed to quantify metabolite overlap within lung tissue positions and between lung tissue and plasma using Intervene Shiny App (45).

Global Natural Products Social Molecular Networking (GNPS) was used to perform feature-based molecular networking and MolNetEnhancer according to the parameters in Table 4. Cytoscape 3.8.2. was used to visualize all molecular networks. All reported annotations are at Metabolomics Standards Initiative confidence level 2 (specific metabolite name provided) or level 3 (metabolite family name provided only) (46). Lipids were annotated based on GNPS library or analog matches, and using standard LipidMAPS nomenclature (47).

**Table 4:** GNPS Parameters

| Feature-Based Molecular Network |  |
| --- | --- |
| Precursor Ion Mass Tolerance | 0.02 |
| Fragment Ion Mass Tolerance | 0.02 |
| Minimum Matched Fragment Ions | 4 |
| Maximum Connected Component Size (Beta) | 100 |
| Maximum shift between precursors | 500 |
| Library Search Min Matched Peaks | 4 |
| Search analogs | Do Search |
| Top results to report per query | 1 |
| Score Threshold | 0.7 |
| Maximum analog difference | 100.0 Da |
| Minimum Peak Intensity | 0.0 |
| Filter Precursor Window | Filter |
| Filter peaks in 50 Da Window | Filter |
| Filter Library | Filter Library |
| Normalization per file | Row Sum Normalization (Per file Sum to 1,000,000) |
| Aggregation Method for peak abundances per group | Mean |
| PCoA | braycurtis |
| Run Dereplicator | Run |

## Data availability

All metabolomics data is publicly available in MassIVE under accession number MSV000085389 (ftp://massive.ucsd.edu/MSV000085389). MOLNet Enhancer link: https://gnps.ucsd.edu/ProteoSAFe/status.jsp?task=f9f194c9b723409f8927c84470f9a0c5 Original GNPS link: https://gnps.ucsd.edu/ProteoSAFe/status.jsp?task=bb7a5f7fb32045208b6040908a1453f7

## Acknowledgments

This project was supported by a pilot grant from the Oklahoma Center for Respiratory and Infectious Diseases (OCRID) under NIH award number P20GM103648 and used the OCRID Animal Model Core, supported by the National Institute of General Medical Sciences under award number P20GM103648. The authors further wish to acknowledge partial support from NIH award number R21AI148886 and to thank Dr. Peter Palese, Icahn School of Medicine at Mount Sinai, Department of Microbiology, New York, New York, USA, for providing the PR8-Glu viral strain. Laura-Isobel McCall, Ph.D. holds an Investigators in the Pathogenesis of Infectious Disease Award from the Burroughs Wellcome Fund. The content is solely the responsibility of the authors and does not necessarily represent the official views of the National Institutes of Health or any of the other funders.

