## Supplementary figures and images for "Spatial metabolomics reveals localized impact of influenza virus infection on the lung tissue metabolome"

### Figure S1

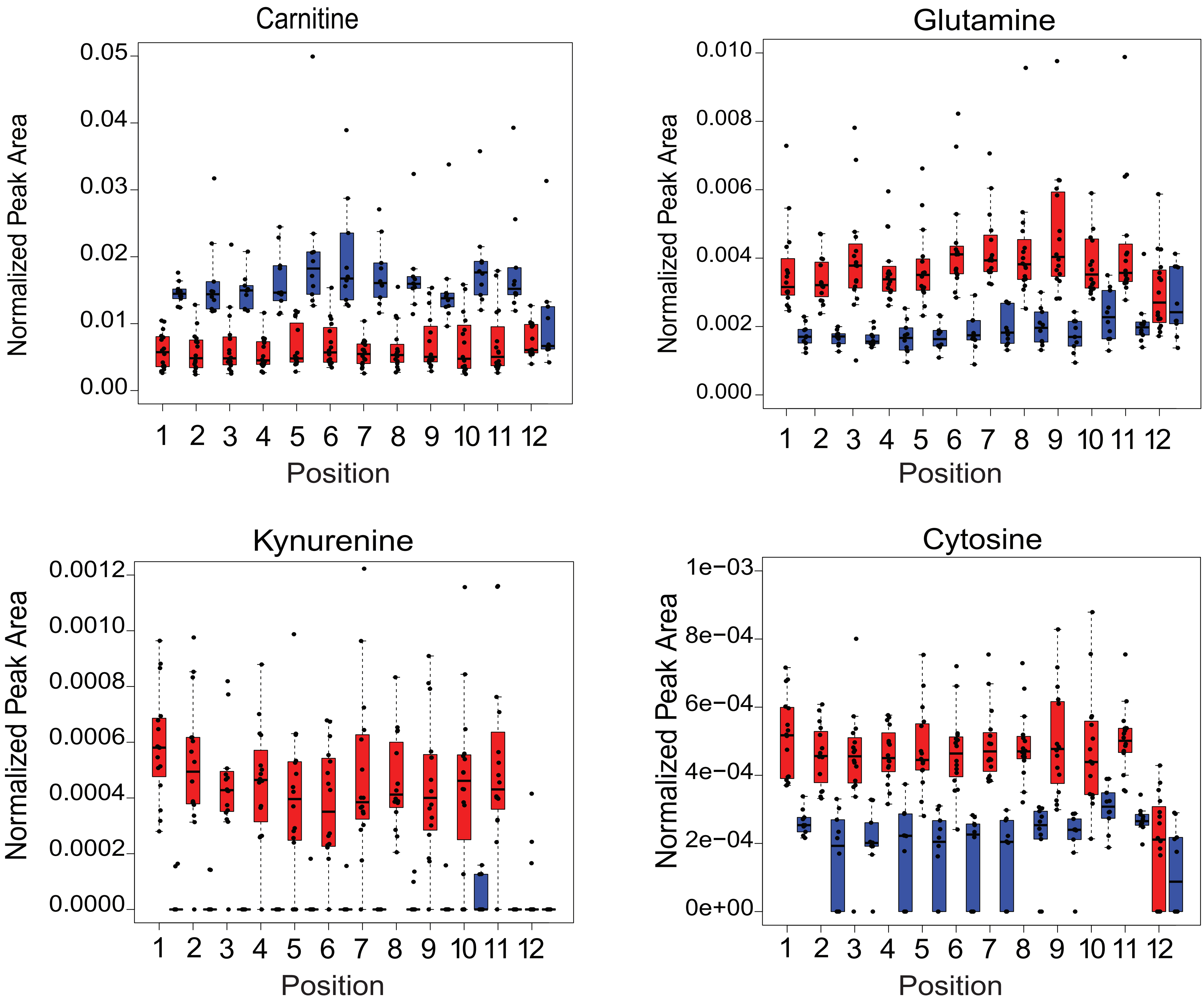

### Figure S2

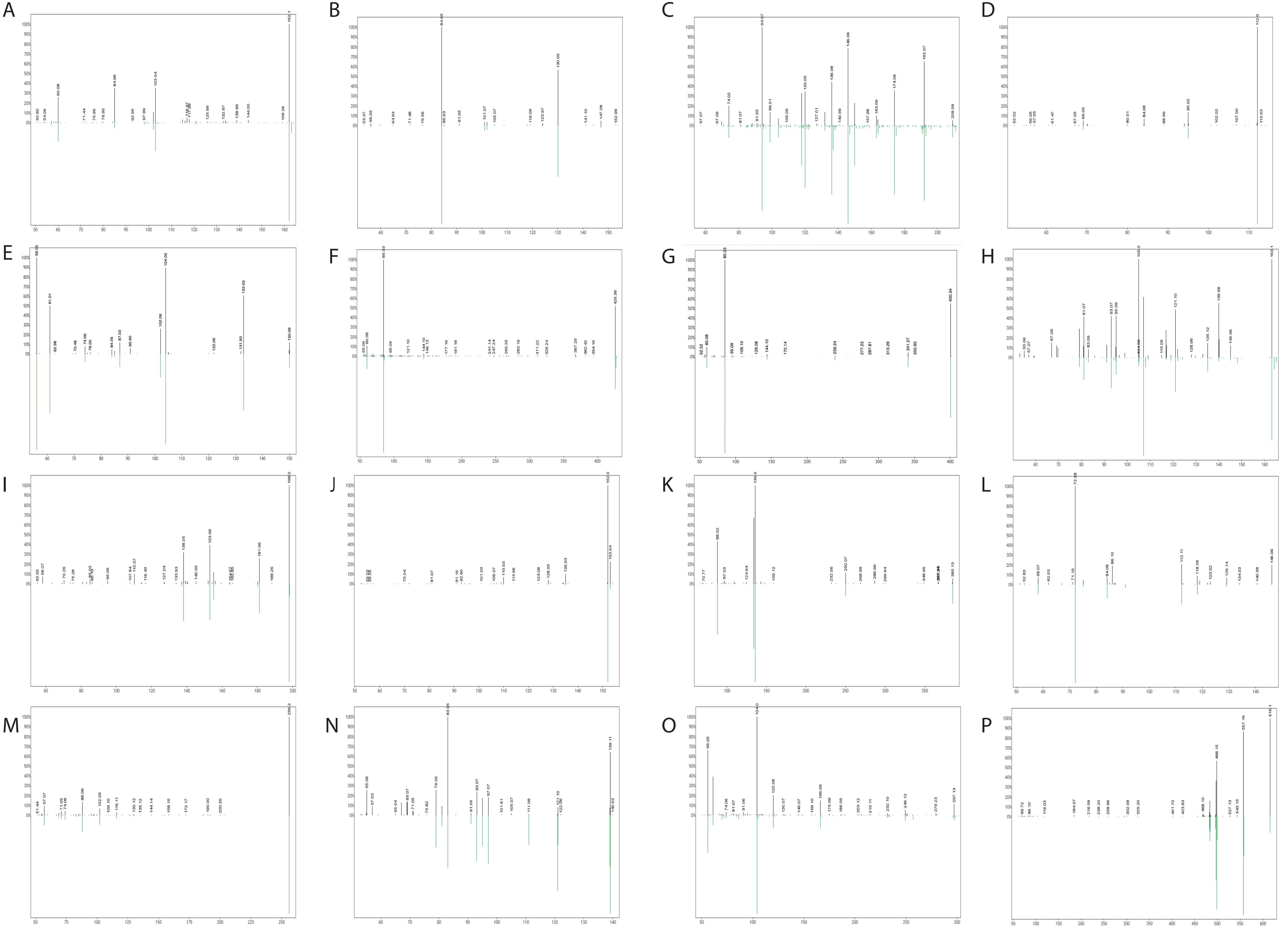

### Figure S3

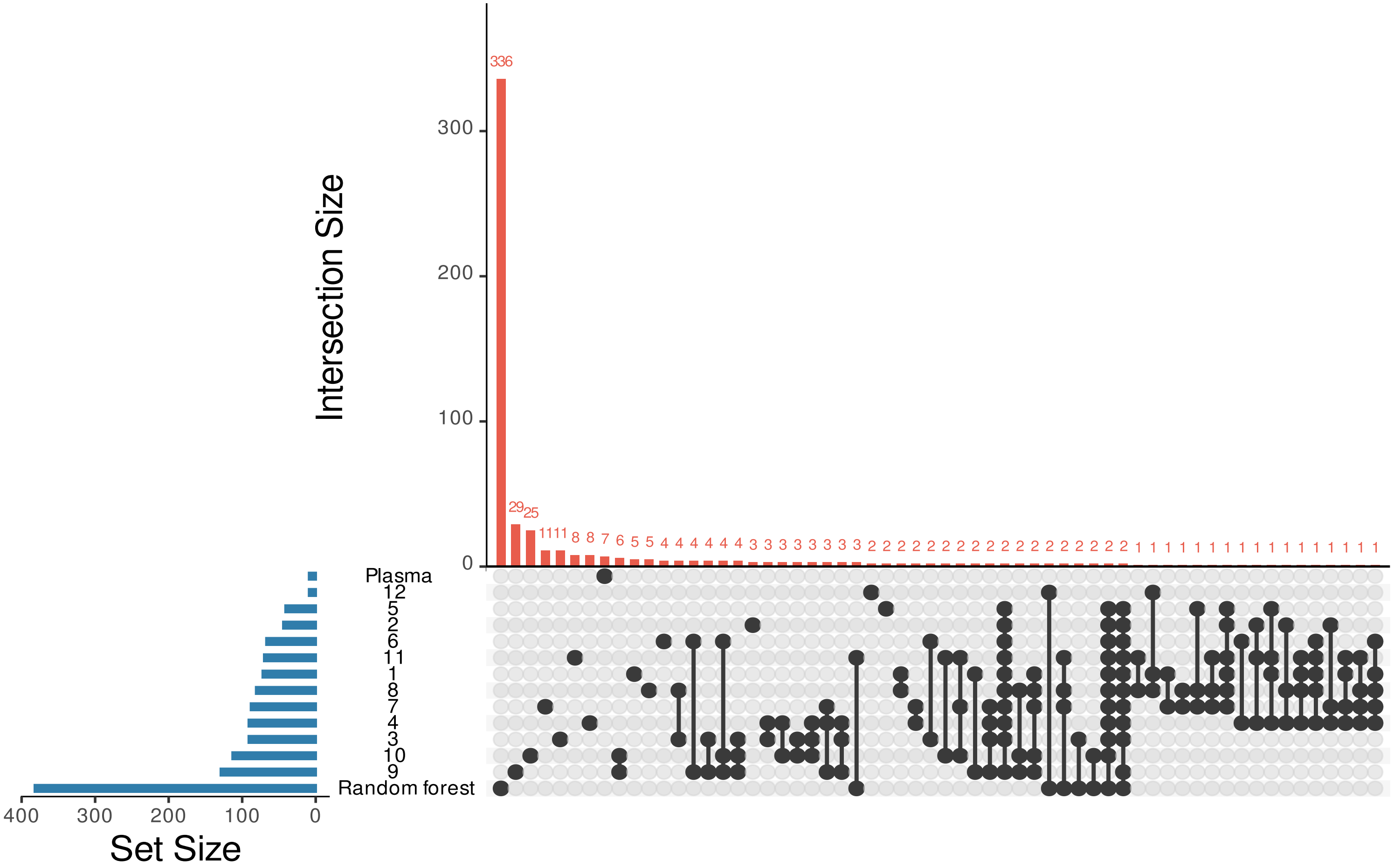
