## Supplementary table and figure legends for "Spatial metabolomics reveals localized impact of influenza virus infection on the lung tissue metabolome"

**Supplementary Information**

**Table S1: Infection-perturbed metabolites in the lung (all positions combined).** NA, not applicable.

**Table S2: Infection-perturbed metabolites in the plasma.** NA, not applicable.

**Table S3: Infection-perturbed metabolites in the lung at position 1.** NA, not applicable.

**Table S4: Infection-perturbed metabolites in the lung at position 2.** NA, not applicable.

**Table S5: Infection-perturbed metabolites in the lung at position 3.** NA, not applicable.

**Table S6: Infection-perturbed metabolites in the lung at position 4.** NA, not applicable.

**Table S7: Infection-perturbed metabolites in the lung at position 5.** NA, not applicable.

**Table S8: Infection-perturbed metabolites in the lung at position 6.** NA, not applicable.

**Table S9: Infection-perturbed metabolites in the lung at position 7.** NA, not applicable.

**Table S10: Infection-perturbed metabolites in the lung at position 8.** NA, not applicable.

**Table S11: Infection-perturbed metabolites in the lung at position 9.** NA, not applicable.

**Table S12: Infection-perturbed metabolites in the lung at position 10.** NA, not applicable.

**Table S13: Infection-perturbed metabolites in the lung at position 11.** NA, not applicable.

**Table S14: Infection-perturbed metabolites in the lung at position 12.** NA, not applicable.

**Table S15: $R^2$ of metabolic perturbation in lung tissue**

**Fig S1: Boxplots of metabolites perturbed by IAV infection in lung tissue (Red - infected, Blue - Uninfected).** Carnitine statistically different at all positions (p-value <0.05). Glutamine statistically different at all positions (p-value <0.05). Kynurenine statistically different at all positions (p-value <0.05). Cytosine statistically different at all positions (p-value <0.05).

**Fig S2: Representative mirror plots of metabolites perturbed by IAV infection in the lung tissue and plasma. A:** Mirror plot of *m/z* 162.113, RT 0.297 min (top, black) to reference library spectrum (L-carnitine, bottom, green). **B:** Mirror plot of *m/z* 147.076, RT 0.32 min (top, black) to reference library spectrum (Glutamine, bottom, green. **C:** Mirror plot of *m/z* 209.092, RT 0.631 min ( top, black) to library spectrum (Kynurenine, bottom, green). **D:** Mirror plot of *m/z* 112.0508, RT 0.31 min (top, black) to reference library spectrum (Cytosine, bottom, green). **E:** Mirror plot of *m/z* 159.058, RT 0.323 min (top, black) to reference library spectrum (Methionine, bottom, green). **F:** Mirror plot of *m/z* 426.357, RT 2.982 min (top, black) to reference library spectrum (Oleyl L-carnitine, buttom, green). **G:** Mirror plot of *m/z* 400.341, RT 2.952 min (top, black) to reference library spectrum (Palmitoylcarnitine, bottom, green). **H:** Mirror plot of *m/z* 166.042, RT 2.246 min (top, black) to reference library spectrum (Phenylalanine, bottom, green). **I:** Mirror plot of *m/z* 198.085, RT 0.309 min (top, black) to reference library spectrum (L-citrulline, bottom, green). **J:** Mirror plot of *m/z* 150.977, RT 0.291 min (top, black) to reference library spectrum (Guanine, bottom, green). **K:** Mirror plot of *m/z* 385.212, RT 2.166 min (top, black) to reference library spectrum (S-Adenosyl-L-homocysteine, bottom, green). **L:** Mirror plot of *m/z* 146.165, RT 0.287 min (top, black) to reference library spectrum (Spermidine, bottom, green). **M:** Mirror plot of *m/z* 256.073, RT 2.563 min (top, black) to reference library spectrum (Palmitamide, bottom, green). **N:** Mirror plot of *m/z* 139.112, RT 2.644 min (top, black) to reference library spectrum (4-Hydroxynonenal, bottom, green). **O:** Mirror plot of *m/z* 297.123, RT 2.444 min (top, black) to reference library spectrum (Met-Phe, bottom, green). **P:** Mirror plot of *m/z* 616.358, 2.994 min (top, black) to reference library spectrum (Hemin cation, bottom, green).

**Fig S3: Limited overlap of metabolites differing in abundance between plasma and lung tissue and metabolites impacted by infection in plasma and at each lung position.** Metabolites differing between plasma and lung tissue were identified using random forest, with a cutoff of >1 ("Random forest" set). Likewise, metabolites differing between infected and uninfected samples were identified by random forest at each tissue site and in plasma individually, as in Fig. 2A (sets 1-12, representing lung segments 1-12, and plasma).
